# Profiling endogenous airway proteases and antiproteases and measuring proteolytic activation of Influenza HA using *in vitro* and *ex vivo* human airway surface liquid samples

**DOI:** 10.1101/2024.06.14.599031

**Authors:** Stephanie A Brocke, Boris Reidel, Camille Ehre, Meghan E Rebuli, Carole Robinette, Kevin D Schichlein, Christian A Brooks, Ilona Jaspers

## Abstract

Imbalance of airway proteases and antiproteases has been implicated in diseases such as COPD and environmental exposures including cigarette smoke and ozone. To initiate infection, endogenous proteases are commandeered by respiratory viruses upon encountering the airway epithelium. The airway proteolytic environment likely contains redundant antiproteases and proteases with diverse catalytic mechanisms, however a proteomic profile of these enzymes and inhibitors in airway samples has not been reported. The objective of this study was to first profile extracellular proteases and antiproteases using human airway epithelial cell cultures and *ex vivo* nasal epithelial lining fluid (NELF) samples. Secondly, we present an optimized method for probing the proteolytic environment of airway surface liquid samples (*in vitro* and *ex vivo*) using fluorogenic peptides modeling the cleavage sites of respiratory viruses. We detected 48 proteases in the apical wash of cultured human nasal epithelial cells (HNECs) (n=6) and 57 in NELF (n=13) samples from healthy human subjects using mass-spectrometry based proteomics. Additionally, we detected 29 and 48 antiproteases in the HNEC apical washes and NELF, respectively. We observed large interindividual variability in rate of cleavage of an Influenza H1 peptide in the *ex vivo* clinical samples. Since protease and antiprotease levels have been found to be altered in the airways of smokers, we compared proteolytic cleavage in *ex vivo* nasal lavage samples from male/female smokers and non-smokers. There was a statistically significant increase in proteolysis of Influenza H1 in NLF from male smokers compared to female smokers. Furthermore, we measured cleavage of the S1/S2 site of SARS-CoV, SARS-CoV-2, and SARS-CoV-2 Delta peptides in various airway samples, suggesting the method could be used for other viruses of public health relevance. This assay presents a direct and efficient method of evaluating the proteolytic environment of human airway samples in assessment of therapeutic treatment, exposure, or underlying disease.

## Introduction

The human genome encodes 703 enzymes with known proteolytic activity and 1,652 endogenous antiproteases (1) which combined makes up over 10% of all protein coding genes in humans. Proteases and their endogenous antiproteases are fundamental to cellular homeostasis (2) and are expressed intracellularly, as membrane-bound proteins, and secreted extracellularly (3). The mucosal surface of the airway represents a complex proteolytic landscape which must interact with environmental pathogens, allergens, and pollutants. There is evidence that the epithelial surface expresses a multitude of proteases with diverse catalytic mechanisms (4–8), but to our knowledge few studies have profiled proteases in the airway (9, 10), and none have specifically profiled antiproteases.

Proteases and antiproteases are important for cell- and tissue-level homeostasis and physiologic function. In the airways, a balance of proteases and antiproteases is necessary for bronchoconstriction (11–13), regulation of mucins and airway surface liquid (14–16), cell differentiation (17), development (18), repair, regeneration (19, 20), and immunity (21, 22). Therefore, many pulmonary pathologies result from aberrant protease elevation or antiprotease depletion which perturb this complex proteolytic activity balance. The role of proteases in manifestation of respiratory diseases has been reviewed previously (23, 24).

Increased susceptibility to viral infection is one outcome of aberrant proteolytic activity in the respiratory tract (25). To initiate infection, viral fusion proteins such as hemagglutinin (HA) in Influenza viruses and the Spike (S) protein in coronaviruses must be cleaved. This cleavage triggers activation of the viral fusogenic machinery, enabling fusion of the host and viral membranes and deposition of genetic material into the cytosol of the host cell. Respiratory viruses have evolved to commandeer extracellular and membrane-bound proteases expressed along the respiratory tract to cleave their fusion proteins and activate infection (26–28). The HA subtypes in strains of mammalian Influenza A which continue to circulate in the human population, including H1, H2 and H3, are cleaved by a number of airway proteases: human airway trypsin-like protease (HAT) (29) , matriptase (30, 31), kallikrein-related proteases (32), transmembrane serine protease 2 (TMPRSS2) (29, 33, 34), and plasmin (35). The S protein of SARS coronaviruses is also cleaved by some of the same proteins (36–39). Because of the role these proteases play in pathogenesis of viral infection, the therapeutic use of antiproteases as a prophylactic measure against infection has been extensively investigated (36, 40–44). Furthermore, environmental exposures, such as cigarette smoke and ozone, have been found to alter secretion of proteases and antiproteases in the airways (45–47). However, the effect these alterations have on susceptibility to viral infections remains understudied.

Proteases in the airway overlap in their substrate specificities, as exemplified above, and this is especially true within enzyme clans, which are designated based on mechanism of catalysis. Furthermore, endogenous inhibitors of proteases do not inhibit proteases of the same clan with equal efficacy (48). Predicting general proteolytic activity of airway surfaces by protein detection or quantification methods is therefore difficult due to the abundance, diversity, and redundancy of proteases and antiproteases expressed there. Evaluation of Influenza HA cleavage by airway proteases has been used to approximate or determine susceptibility to infection using two general methods: (i) use of fluorogenic substrates modeling the Influenza HA cleavage site to investigate cleavage by individual proteases (35, 49), and (ii) use of Western blotting to detect HA cleavage fragments by individual enzymes or cell culture supernatants (31, 46, 50, 51).

In the present study, we sought to first profile the diverse and abundant proteases and antiproteases secreted from *in vitro* primary airway epithelial cells grown at air-liquid interface (ALI) as well as *ex vivo* nasal epithelial lining fluid (NELF) from healthy human donors. Next, we present a novel methodology for probing the proteolytic activity of these samples toward Influenza H1 using a non-infectious internally quenched fluorescent (IQF) peptide modelled after the viral cleavage site (Table 1). We also demonstrate utility of this assay towards additional viral substrates, SARS-CoV-1 S and SARS-CoV-2 S. Using organotypic cultures of nasal and bronchial epithelial cells, we compared proteolytic activation of the viral peptides in multiple airway regions. This method offers a more high-throughput and straightforward approach to assessing susceptibility to Influenza infection relative to immunoassay- or gel-based detection methods which also require larger sample volumes. Comparison of proteolytic activation of viral substrates pre- and post-environmental exposure, in potentially susceptible groups, or as a screen for efficacy of preventative therapeutics represent applications of this method. To this end, we evaluated rates of Influenza H1 cleavage in previously collected nasal lavage fluid (NLF) from smokers and non-smokers.

**Table 1.** Amino acid sequences of peptides used for experimentation and residue numbers from the full-length proteins. For each peptide, the site of cleavage occurs between the two residues highlighted in red. The SARS coronavirus peptides model the S1/S2 cleavage site. The multi-basic Furin cleavage site insertion acquired by SARS-CoV-2 is underlined. MCA = 7-methoxycoumarin-4-yl acetyl, DNP = N-2,4-dinitrophenyl.

| Peptide | Amino acid residues | Sequence (N'-C') | N'-terminus modification | C'-terminus modification | GenBank Accession |
| --- | --- | --- | --- | --- | --- |
| Influenza HA A/California/04/2009 H1N1 | 338-346 | IPSIQSRGL | MCA | Lysine-DNP | ACP41105 |
| SARS-CoV-1 S wild type | 661-671 | HTVSLLRSTSQ | MCA | Lysine-DNP | ABD73002 |
| SARS-CoV-2 S wild type | 678-688 | TNSPRRAR<u>SV</u>A | MCA | Lysine-DNP | QIG55955 |
| SARS-CoV-2 S Delta variant | 678-688 | TNSRRRAR<u>SV</u>A | MCA | Lysine-DNP | UAL04647 |

## Materials and Methods

### Culture of HNECs and HBECs

Primary human nasal epithelial cells (HNECs) were obtained from healthy adult volunteers aged 18-59 years. Demographic information of HNEC donors is provided (Table 2). Subjects were recruited between November 1, 2019 and March 11, 2021 by a protocol approved by the Institutional Review Board at the University of North Carolina (Protocol # IRB 11-1363). Written informed consent was obtained from all subjects. Complete sample collection and culturing methods have been published previously (52), but briefly, inferior turbinate nasal scrapes were collected from each donor and expanded in flasks using PneumaCult-Ex Plus Medium (STEMCELL, Vancouver, BC, Canada) supplemented with 1% penicillin-streptomycin at 5% CO_2_ and 37°C. After two passages, cells were frozen down using Bambanker medium (Lymphotec, Tokyo, Japan) and stored in liquid nitrogen. For experimentation, cells were thawed and seeded in a flask for one additional expansion. Upon confluency, cells were seeded onto permeable polyethylene terephthalate 12-well inserts with a 0.4 μm pore size (CELLTREAT, Pepperell, MA, USA) which were coated with human placental type IV collagen (Sigma-Aldrich, St. Louis, MO; C7521). Ex Plus medium was added to both the apical and basolateral compartments and changed daily until the cells reached confluency on the inserts. At confluency, Ex Plus medium was exchanged for PneumaCult ALI medium (STEMCELL, Vancouver, BC, Canada) on the basolateral side and medium was removed on the apical side. For one week, basolateral medium was replaced daily. From then on, three times per week the medium was changed, and the apical surface was washed with Hanks Balanced Saline Solution +CaCl_2_, +MgCl_2_, (HBSS++). Cells were differentiated at ALI conditions for >30 days, until ciliation and mucus were present on the cultures.

**Table 2.** Demographic information of HNEC donors used for proteomic analysis (n=6) as well as HNEC (n=13) and HBEC (n=3) donors used for proteolytic cleavage assays.

|  | **HNECs for proteomics**<br>**(n=6)** | **HNECs (n=13)** | **HBECs (n=3)** |
| --- | --- | --- | --- |
| Age, mean±standard deviation | 48.5±13.9 | 26.9±6.1 | 36.7±4 |
| Sex, n, M/F | 4/2 | 8/5 | 2/1 |
| Race, n, Black/White/Asian | 0/6/0 | 1/10/2 | 1/2/0 |
| Ethnicity, n, Hispanic/Non-Hispanic | 1/5 | 2/11 | 0/3 |

Primary human bronchial epithelial cells (HBECs) were sourced from non-diseased, non-smoker adult donors through the Marsico Lung Institute Tissue Procurement and Cell Culture Core at the University of North Carolina. Demographic data are provided in Table 2. Cells were cultured and differentiated as described previously (53, 54). The cells were initially cultured in flasks coated with bovine type-1 collagen (Advanced BioMatrix, Carlsbad, CA, USA) in PneumaCult-Ex Plus medium (STEMCELL, Vancouver, BC, Canada) supplemented with 1% penicillin-streptomycin. Upon reaching 70-90% confluence, the cells were passaged with Accutase (Innovative Cell Technologies, San Diego, CA, USA). Passage 3 cells were plated on 12 mm, 0.4-µm permeable cell culture inserts (CELLTREAT, Pepperell, MA, USA) coated with human placental type IV collagen (Sigma-Aldrich, St. Louis, MO, USA). The cells were cultured using Ex Plus in both the basolateral and apical compartments for 1 week. Upon achieving confluence, the cells were subjected to differentiation at an air-liquid interface (ALI) for at least 30 days with Pneumacult ALI (STEMCELL, Vancouver, BC, Canada) growth medium in the basolateral compartment. Basolateral media was initially replaced daily for 1 week, then subsequently replaced three times a week, and the apical surface was washed once a week with HBSS++ to clear apical mucus and cellular debris.

### HNEC and HBEC sample collection

Once cells were differentiated, apical washes for proteolytic cleavage assays were collected by adding 200 μl of 37°C HBSS++ to the apical surface of each culture and incubating at room temperature for 15 minutes. Apical wash liquid was then carefully removed with a pipette and pooled by donor in microcentrifuge tubes. Samples were stored at -80°C.

### Nasal lavage fluid (NLF) sample collection

NLF was collected and processed from healthy adults aged 18-50 years who were either never-smokers or cigarette smokers defined based on self-reporting and usage of smoking diaries, as described previously (55, 56). Samples were collected between August 1, 2014 and March 1, 2016. Exclusion criteria included symptoms of allergic rhinitis, chronic obstructive pulmonary disease, asthma, and use of immunosuppressive drugs such as corticosteroids. Demographic data from these participants are provided in Table 3. The protocol used for NLF sample collection was approved by the University of North Carolina’s Institutional Review Board (Protocol # IRB 13-3454) and written informed consent was obtained from all subjects.

**Table 3.** Demographic information of NLF donors.

|  | All (n=48) | Cigarette smokers (n=24) | Non-smokers (n=24) |
| --- | --- | --- | --- |
| Age, mean±standard deviation | 30.1±6.7 | 30.8±6.4 | 29.4±6.9 |
| Sex, n, M/F | 24/24 | 12/12 | 12/12 |
| Race, n, Black/White/Asian | 16/31/1 | 12/11/1 | 4/20/0 |

### Nasal epithelial lining fluid (NELF) sample collection

NELF was collected as previously described (57) on October 26, 2023 from n=19 donors by a protocol approved by the Institutional Review Board at the University of North Carolina (Protocol # IRB 11-1363) and written informed consent was obtained from each subject. Demographic information for sample donors is provided in Table 4. Briefly, nostrils were sprayed with 0.9% saline solution, then Leukosorb paper (Pall Scientific, Port Washington, NY, USA) cut in strips to fit the nostrils were inserted into both nares. A padded clip was used to clamp the nostrils closed and keep the Leukosorb strips in place for 2 minutes. The Leukosorb strips were removed and stored in 1.5 ml microcentrifuge tubes at -20°C until elution. Eluate from the Leukosorb strips was collected as previously described (57). A 100 µl volume of 1% BSA + 0.05% Triton X-100 in PBS was pipetted onto each strip. The strips were then centrifuged twice at 13,000 rpm for 2 minutes to collect the eluate at the bottom of the tube.

**Table 4.** Demographic information of NELF donors.

|  | **All (n=19)** |
| --- | --- |
| Age, mean±standard deviation | 29.9±6.4 |
| Sex, n, M/F | 10/9 |
| Race, n, Black/White/Asian | 3/13/3 |
| Hispanic/Latino, n, Y/N | 3/16 |

### Mass Spectrometry-based proteomic analysis

HNEC culture apical secretions and nasal strip samples were prepared for label-free proteomics using filter-aided sample preparation (FASP) (58). HNEC apical washes from n=6 donors (4M, 2F) and NELF from n=13 donors (5M, 8F) were used for proteomics. For HNEC secretions 100 µl apical washes were denatured using 8M urea, and nasal strips were extracted in 0.5 ml 4M GuHCl, followed by the reduction of cysteine residues by 10 mM dithiothreitol (Sigma-Aldrich, St. Louis, MO, USA) and alkylation in 50 mM iodoacetamide (Sigma-Aldrich). Samples were digested with pig trypsin (25 ng/µl) overnight at 37°C. The resulting peptide mixtures were vacuum freeze-dried and dissolved in 30 µl of 1% acetonitrile and 0.1% trifluoroacetic acid, and 5 µl were injected into each sample for chromatography tandem mass spectrometry (LC-MS/MS). LC-MS/MS analysis was performed utilizing a Q-Exactive (Thermo Scientific, Waltham, MA, USA) mass spectrometer coupled to an Ultimate 3000 nano HPLC system (Thermo Scientific), and data acquisitions were performed as described previously (59). The raw data were processed and searched against the UniProt protein database (Homo sapiens, November 2023) using Proteome Discoverer 1.4 (Thermo Scientific, Waltham, MA, USA) software. The following parameters were used in the Sequest search engine: 10 ppm mass accuracy for parent ions and 0.02 Da accuracy for fragment ions; 2 missed cleavages were allowed. The carbamidomethyl modification for cysteines was set to fixed, and methionine oxidation was set to variable. Scaffold 5.3.0 (Proteome Software Inc., Portland, OR, USA) was used to validate the MS/MS-based peptide and protein identifications. Peptide identifications were accepted if they had greater than 95.0% probability by the scaffold local FDR algorithm. Protein identifications were accepted if they had greater than 99.0% probability and contained at least 2 identified peptides. Protein probabilities were assigned by the Protein Prophet algorithm (60). Proteins that contained similar peptides and could not be differentiated based on MS/MS analysis alone were grouped to satisfy the principles of parsimony. The resulting protein identification lists were annotated using NCBI annotations and filtered using the Gene Ontology Terms “endopeptidase activity” (GO:0004175) plus “exopeptidase activity” (GO:0008238) for the protease lists, and “peptidase inhibitor activity” (GO:0030414) for the protease inhibitor lists.

### Viral IQF peptide design

The design of the IQF peptides was based on the work by Jaimes and Straus et al. (35, 37). The amino acid sequences of all IQF peptides used in our assays are provided in Table 1. An additional peptide modeled after the SARS-CoV-2 Delta variant was also constructed because of its highly conserved P681R mutation in the cleavage site. This mutation is believed to aid in viral fusion with the host and render the strain more pathogenic than the wild-type strain (61). The peptides were modified to include the fluorophore 7-methoxycoumarin-4-yl acetyl (MCA) on the N-terminus and an additional Lysine residue with N-2,4-dinitrophenyl (DNP) as a quencher on the C-terminus. Custom peptides were ordered from Biomatik (Kitchener, Ontario, Canada) and arrived as 1 mg aliquots of lyophilized powder and stored at -20°C. Peptides were ordered with the following specifications; purity >95%, TFA removed, and switched to HCl salt. Before use, the peptides were brought to room temperature then resuspended in pure dimethyl sulfoxide (DMSO) at 1 mg/ml. This concentrated peptide stock was then aliquoted and stored at -80°C.

### Proteolytic cleavage assay protocol

Assays were performed in half-area black 96-well plates (Corning Inc, Corning, NY, USA) with a total reaction volume of 50 μl. Airway culture samples and *ex vivo* airway samples were thawed on ice, vortexed briefly and centrifuged at 8,000 x g for 2 minutes to collect mucus and debris. For each assay, 25 μl of sample was used unless otherwise indicated. The frozen peptide stock aliquots were thawed and diluted to 250 μM in PBS with 25% DMSO. Then 10 μl of diluted peptide was added to each well for a final assay peptide concentration of 50 μM, 5% DMSO. PBS was added for its buffering capacity to bring the total assay volume to 50 μl. A multichannel pipet was used to add the dilute peptide to the assay plate immediately prior to assay initiation to increase substrate loading efficiency. A mixture of protease inhibitors was used when indicated, which consisted of 33 μM camostat mesylate (a serine protease inhibitor (62)) (Bio-Techne, Minneapolis, MN, USA), 33 μM decanoyl-RVKR-Chloromethylketone (a proprotein convertase inhibitor (63)) (Bio-Techne, Minneapolis, MN, USA), and 33 μM E64 (a cysteine protease inhibitor (64)) (Sigma-Aldrich, St. Louis, MO, USA). Additionally, recombinant human Furin (New England Biolabs Inc., Ipswich, MA, USA) was diluted in PBS and added to assays as indicated at a concentration of 10 U/ml. Fluorescence was measured with an excitation wavelength of 330 nm and emission wavelength of 390 nm using a CLARIOstar Plus plate reader (BMG Labtech, Ortenburg, Germany). Fluorescence intensity from each well was averaged across 16 locations in the well and was measured approximately once every 1-2 minutes (dependent upon total cycle length) for 60 cycles. The gain setting remained constant for all assays. For all data shown, maximum of slope measurements were based on a width of 6 minutes.

### Proteolytic cleavage assay on ALI culture surface

Proteolytic activity was measured directly on the apical surface of two replicate cultures from n=3 HBEC donors. Immediately prior to the assay, 100 μl of assay solution with 50 μM peptide and 5% DMSO in PBS were pipetted onto the apical surface of the cultures growing on CELLTREAT culture inserts. The plate was then read in a CLARIOstar plate reader as described above, but an Atmospheric Control Unit was additionally used to maintain the assay chamber at 37°C and 5% CO_2_ for the duration of the assay to maintain culture quality. Assays were conducted in clear, 12-well cell culture plates (Corning Inc, Corning, NY, USA) with inserts in place and 1 ml of Pneumacult ALI medium in the basolateral compartment.

### Statistical analysis

Statistical tests and data visualization were performed using GraphPad Prism software (v 10.2.0). ANOVA and *t* tests were used to test for differences between groups and paired/repeated measures versions of the tests were implemented where appropriate. Bonferroni’s post hoc test was used in combination with ANOVA. P values ≤ 0.05 were considered statistically significant.

## Results

### Proteomic profiling of proteases and antiproteases in human airway samples

The landscape of proteases and antiproteases present in HNEC apical washes and NELF samples from healthy donors was evaluated with mass spectrometry-based proteomics. The proteases detected uniquely in each sample type as well as in both sample types are shown in Table 5, likewise with antiproteases in Table 6. A greater number of proteases and antiproteases were detectable in NELF samples (57 and 48 respectively) compared to HNEC apical wash (48 and 29). There were 26 proteases and 19 antiproteases present in both sample types. Enzymes with a variety of catalytic mechanisms were detected, including cysteine proteases, serine proteases, and metalloproteases. Accordingly, protease inhibitors against multiple catalytic mechanisms were also detected; cystatins (cysteine protease inhibitors), SERPINs (serine protease inhibitors), TIMPs (tissue inhibitors of metallopeptidases), and others.

**Table 5.**
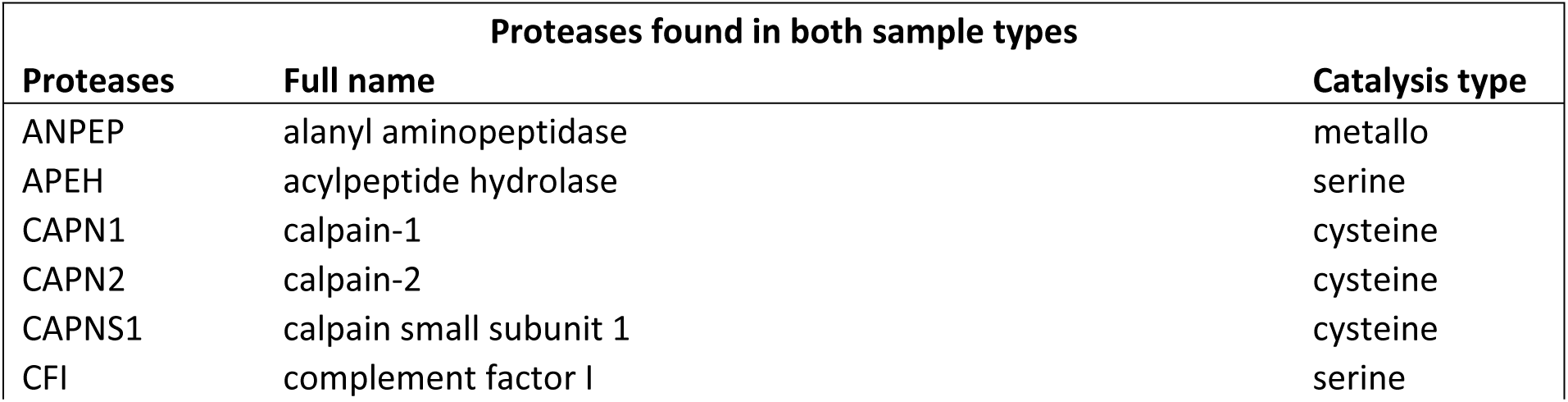
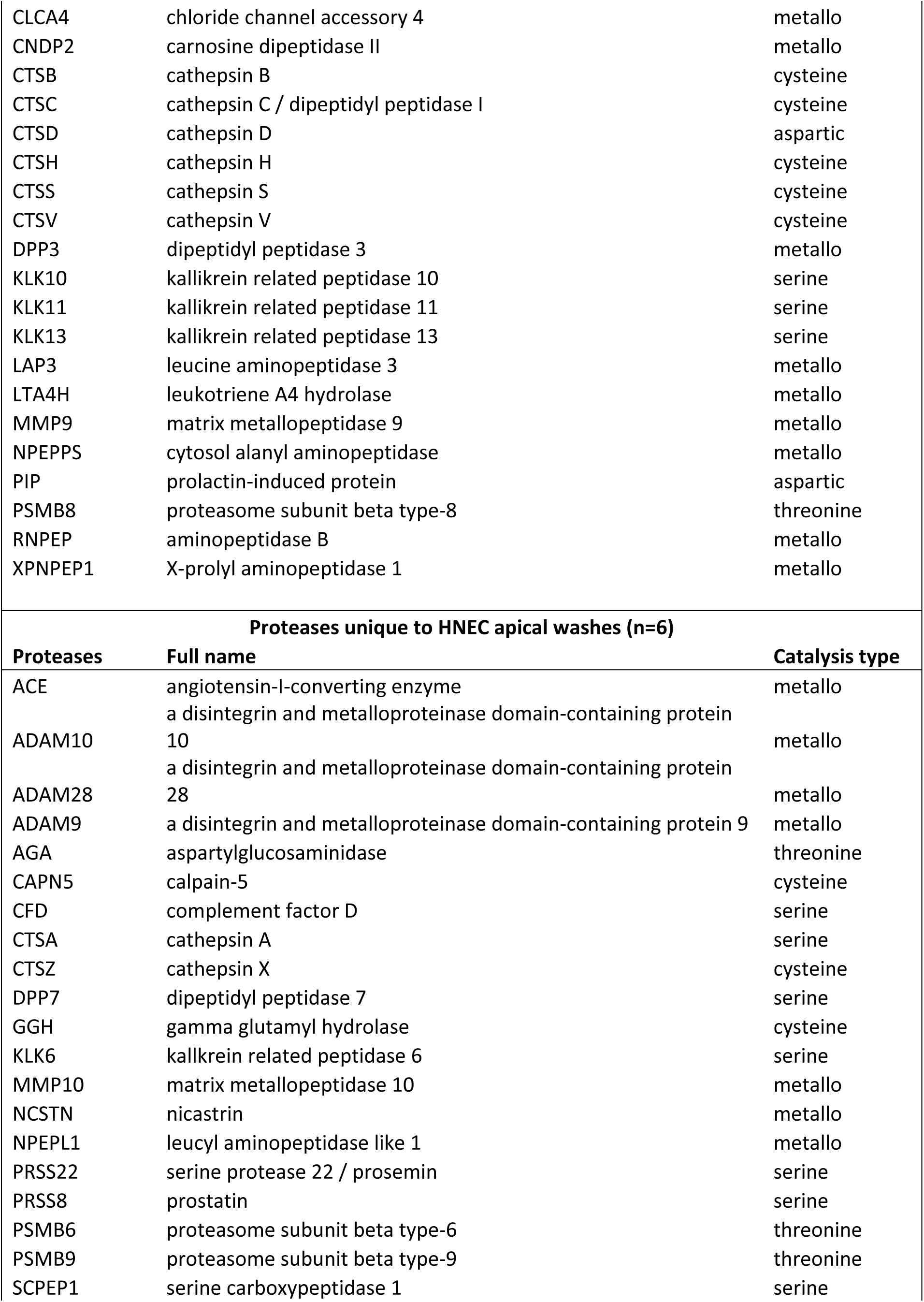
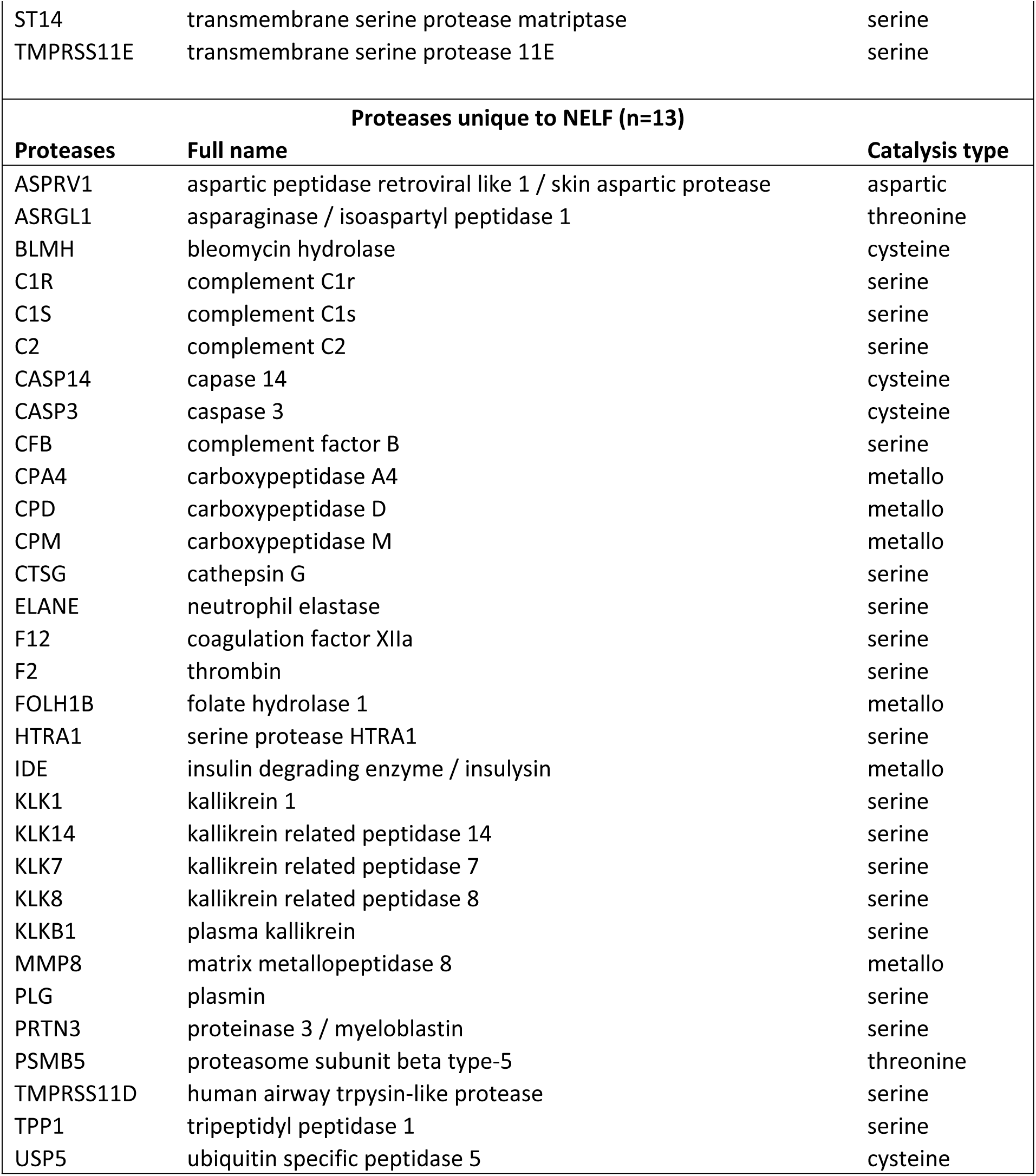
List of proteases detected in both sample types as well as proteases found uniquely in HNEC apical wash samples (n=6) and NELF samples (n=13) by mass spectrometry based proteomic analysis.

**Table 6.**
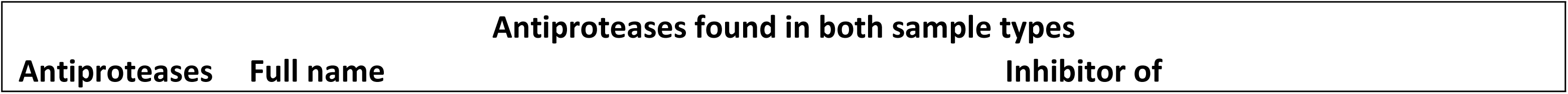
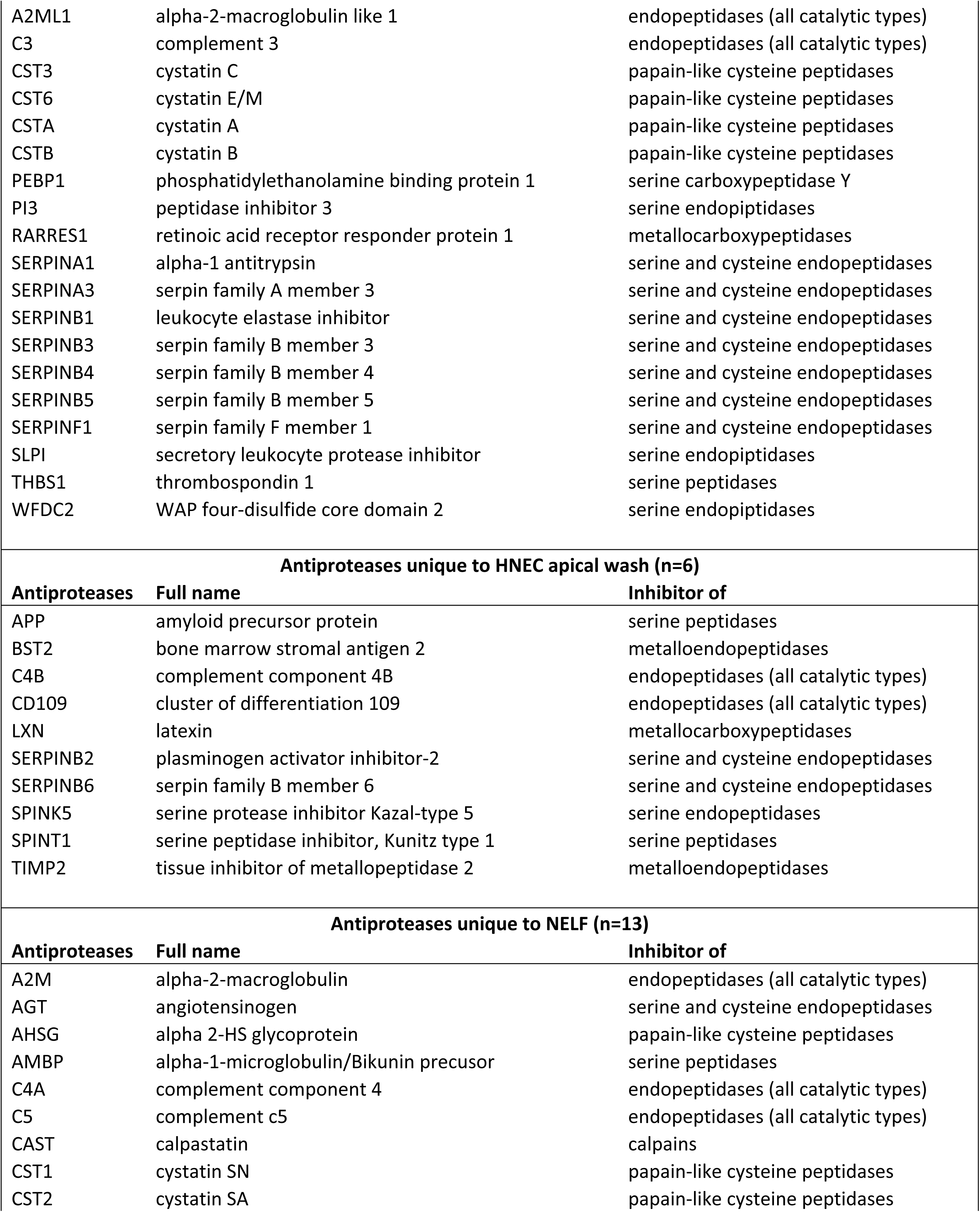
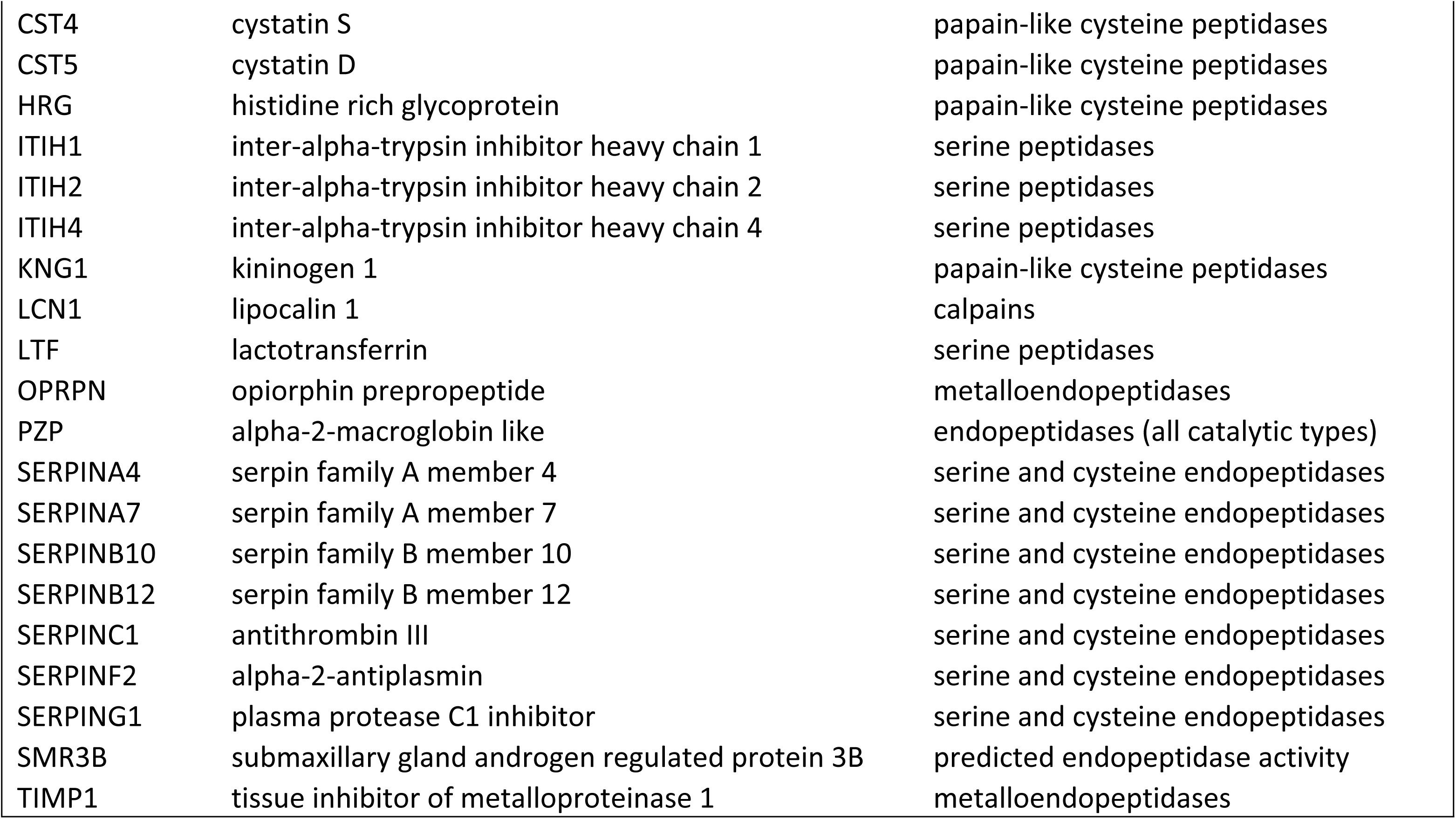
List of antiproteases detected in both sample types as well as uniquely in HNEC apical wash samples (n=6) and NELF samples (n=13) by mass spectrometry based proteomic analysis.

### Measuring protease activity in HNEC mucociliary surface secretions

Apical secretions in differentiated HNEC from n=9 donors were tested for proteolytic activity toward the Influenza H1 IQF peptide. Combining apical wash liquid with the IQF peptides resulted in an increase in fluorescence intensity over time relative to assaying the peptide alone at the same concentration (50 μM), shown in Figure 1A. Addition of a mixture of protease inhibitors against serine, cysteine, and Furin-like proteases resulted in a statistically significant reduction in the maximum slope of the cleavage reaction (change in fluorescence intensity versus time) of Influenza H1 (Figure 1B). Of note, there was a statistically significant difference in age of HNEC donors used for proteomics versus for proteolytic cleavage assays (p=0.0419) using Brown-Forsythe and Welch’s ANOVA tests, with Dunnett’s post hoc test. For proteomic analysis, the mean age of HNEC donors was 48.5±13.9 years and the mean age of HNEC donors for proteolytic cleavage assays was 26.9±6.1 years.

**Fig 1.**
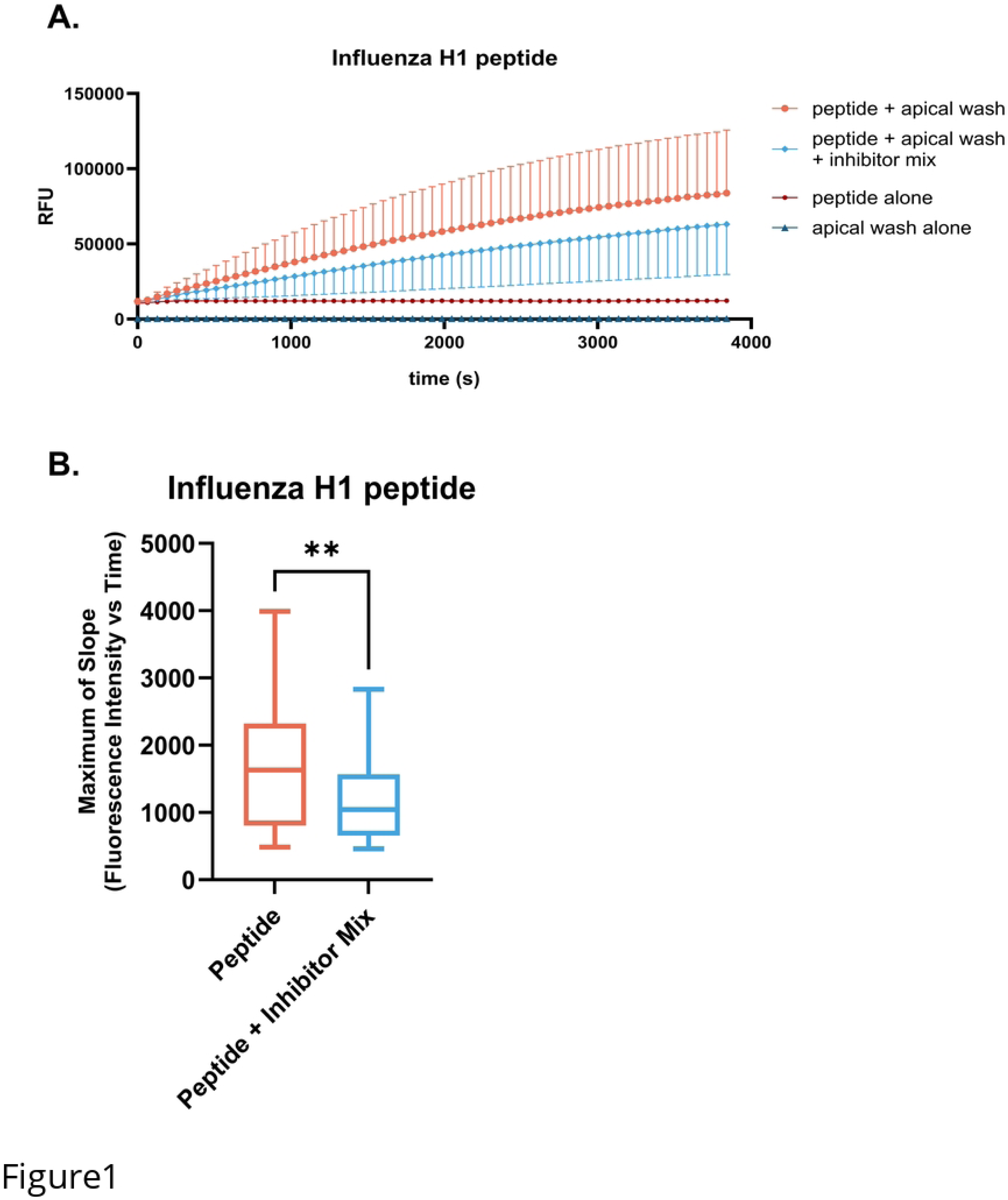
Proteases in HNEC apical wash cleave Influenza H1. Cleavage of the IQF Influenza H1 peptide was tested using apical wash samples from n=9 HNEC donors. A) Change in relative fluorescence units (RFU) versus time (s). Addition of a protease inhibitor mix on rate of cleavage is also shown. Samples were collected 4 d since the prior apical wash. B) Maximum of slope of the cleavage reaction is plotted and a paired *t*-test was used to evaluate differences between groups; ** indicates p≤0.01.

### Proteases accumulate on the apical surface of ALI cultures over time

After an initial wash at time=0, apical washing with HBSS++ was repeated on separate cultures from multiple HNEC donors at 24, 48, 72, or 96h. Figure 2 demonstrates that proteolytic activity of HNEC apical secretions toward Influenza H1 increased with duration since the prior apical wash, albeit the change was not statistically significant. Furthermore, apical washes from n=5 HNEC donors demonstrated stable proteolytic activity toward the Influenza H1 IQF peptide from zero to four freeze and thaw cycles (Supplemental Figure 1), suggesting feasibility of applying the proteolytic cleavage assay to stored samples.

**Fig 2.**
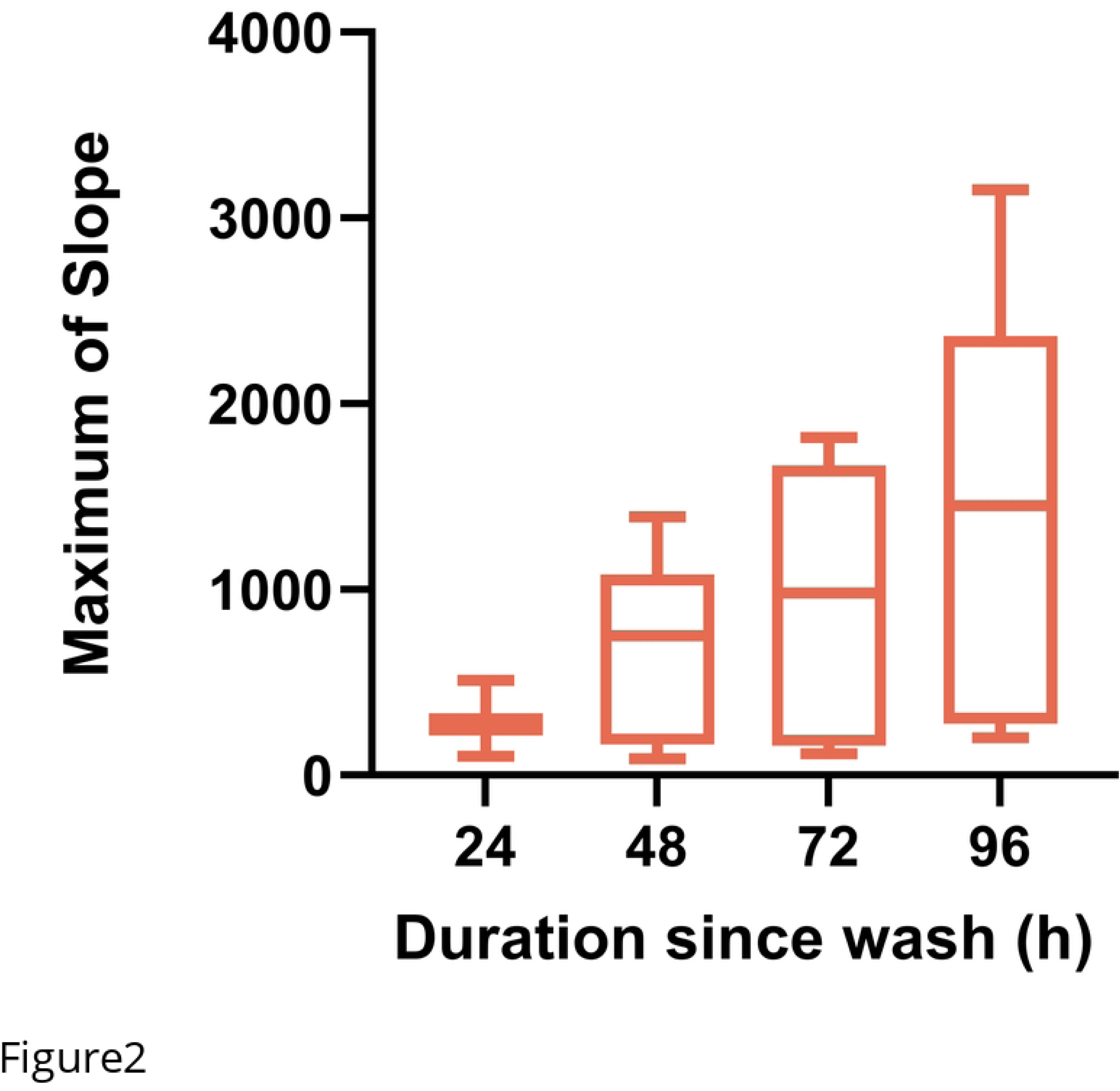
Proteases accumulate on the apical surface of ALI cultures over time. The apical surface of HNEC cultures from n=5 donors was washed at 24, 28, 72, or 96 h post the initial wash.

### Maximum rate of cleavage increases with substrate concentration

The impact of the Influenza H1 substrate concentration on the maximum slope of the cleavage reaction was evaluated. Maximum rates of cleavage with 1, 5, 25, 50, and 500 μM peptide concentrations were tested using apical washes from n=3 HNEC donors. Maximum slope of the cleavage reaction increased with concentration up to 50 μM, and dropped off substantially at 500 μM (Figure 3), suggesting optimal saturation of the assay around 50 μM.

**Fig 3.**
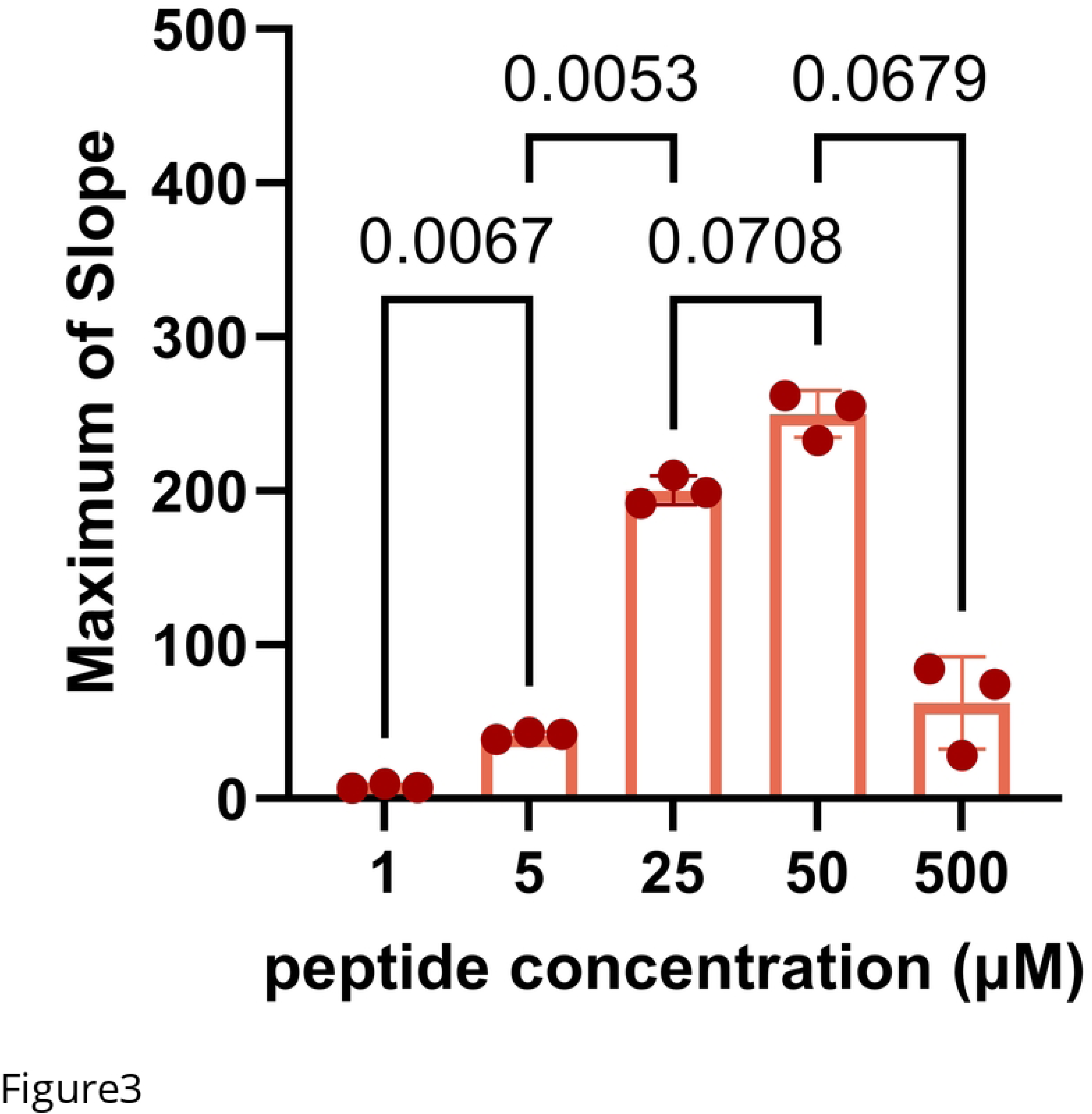
Rate of cleavage increases with substrate concentration up to 50 µM. Maximum rates of cleavage of Influenza H1 peptide at a range of concentrations from 1-500 μM tested with apical wash samples from n=3 HNEC donors. Repeated measures one-way ANOVA with Bonferroni’s post hoc test was used to determine differences between groups, p≤0.05.

### Proteolytic activity toward Influenza H1 is detectable in multiple airway cell types

Like HNECs, human bronchial epithelial cells (HBECs) are also commonly grown at air-liquid interface as an organotypic airway model system, and Influenza has been shown to replicate successfully in both the nasal and bronchial epithelium (65). Proteolytic activity toward Influenza H1 was compared in apical washes from n=5 HNEC and n=3 HBEC donors, which were normalized to a total protein concentration of 58 ng/μl, shown in Figure 4. No statistically significant difference in cleavage of Influenza H1 was observed between the two airway cell types.

**Fig 4.**
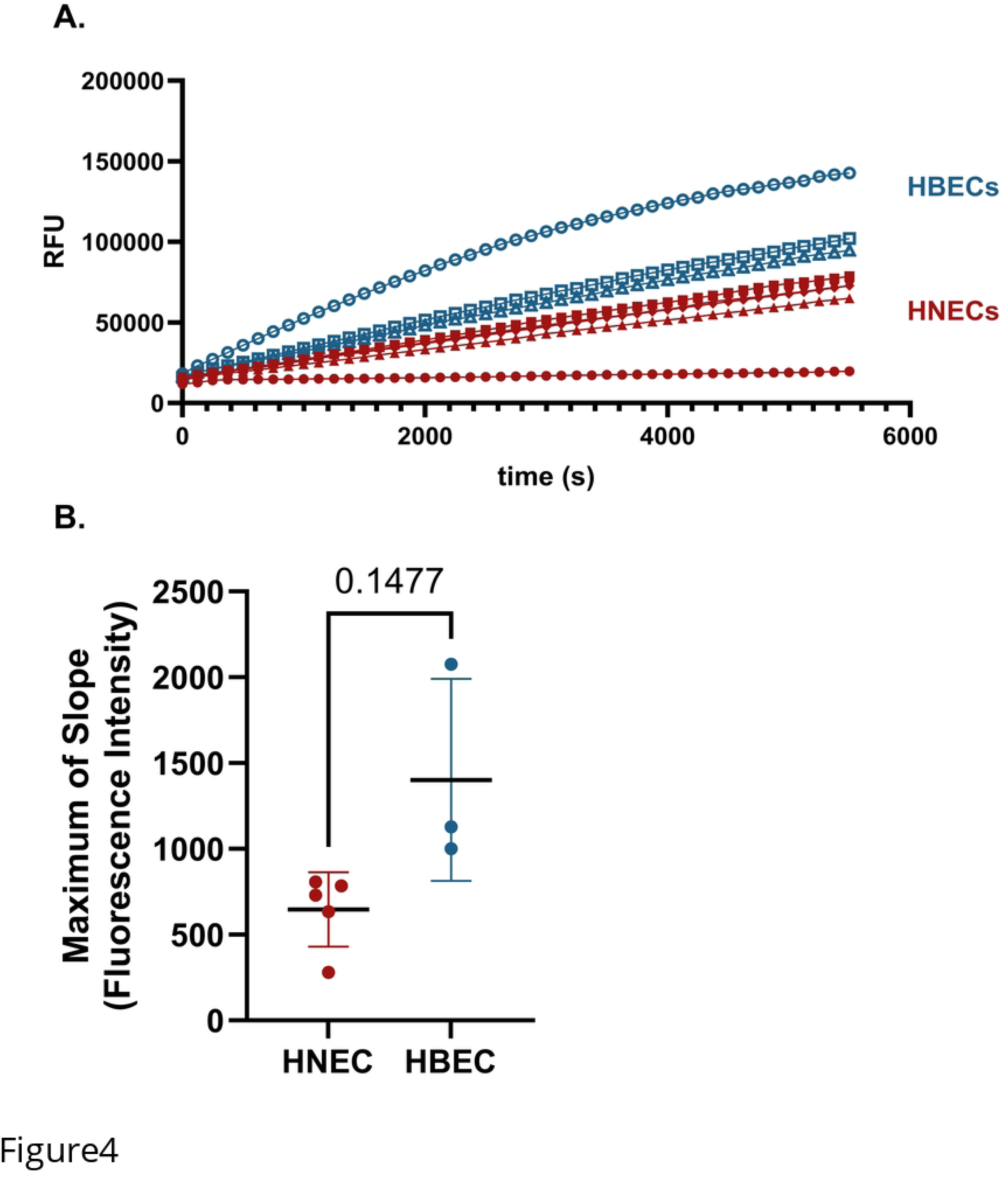
Comparison of Influenza H1 cleavage by proteases in nasal vs bronchial samples. A) Rate of cleavage of Influenza H1 by proteases in apical wash samples from n=5 HNEC donors and n=3 HBEC donors. All samples were collected 7 d from the prior apical wash. Each line represents the average of three technical replicates with standard deviation. B) Maximum of slope of the cleavage reaction is plotted. A Welch’s unequal variances *t*-test was used to test for difference between groups.

### The apical surface of ALI cultures is proteolytically active

While the prior data were generated using apical washes, we also measured proteolytic activity directly on the surface of primary human epithelial cell cultures at air-liquid interface. Cleavage of the Influenza H1 peptide was measured on the apical surface of HBECs (n=3), demonstrating interindividual variability in rates of cleavage between donors (Figure 5).

**Fig 5.**
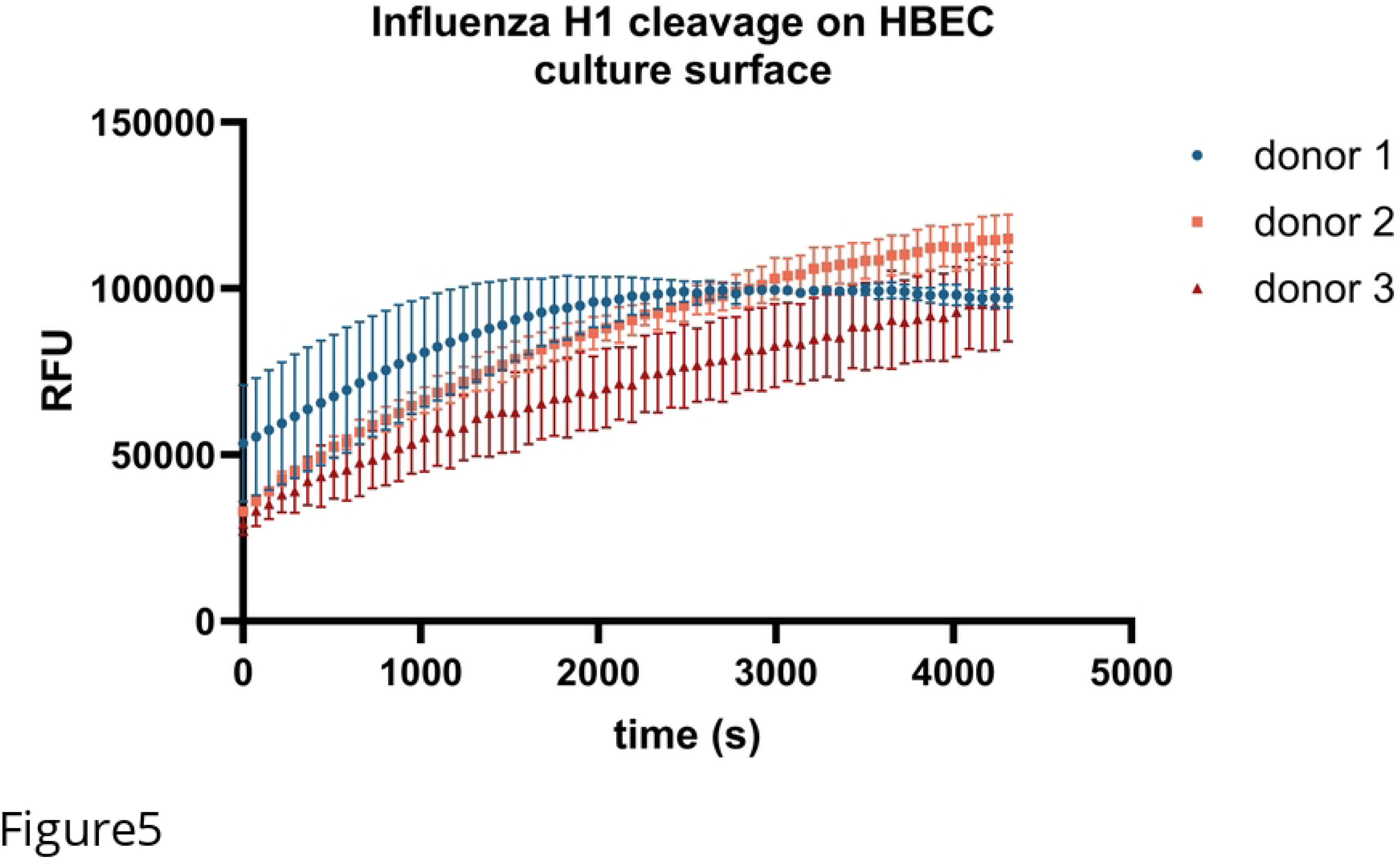
Proteolytic activity measured directly on the apical surface of ALI cultures. Proteolytic activity was measured on HBEC cultures at ALI. A 50 μM solution of Influenza H1 peptide in 100 μl of total volume was added to the apical surface of duplicate cultures from n=3 donors. The prior apical wash of the cultures was 72h prior. Change in fluorescence intensity over time (s) is shown.

### Proteolytic activity toward Influenza H1 can be measured in NLF and NELF

To evaluate whether proteolytic activity towards Influenza virus from clinical samples could be detected by this assay *ex vivo*, we measured rates of cleavage of the viral peptides by nasal lavage fluid samples collected from male and female smokers and non-smokers. As shown in Figure 6, the rate of cleavage of Influenza H1 peptide was higher in males compared to females in the aggregate dataset with both smokers and non-smokers. Stratification by smoking status revealed that cleavage of Influenza H1 was specifically elevated in male smokers. There was no difference in rate of cleavage of Influenza H1 between non-smoking males and females and no difference between smokers and non-smokers. Similarly, we sought to demonstrate proteolytic capacity toward viral substrates in NELF. Rates of cleavage of Influenza H1 was measured using NELF samples from male (n=10) and female (n=9) non-smoking donors. Change in fluorescence intensity versus time for each donor is plotted in Figure 7A. The maximum slope of the cleavage reactions by sex of donor is also shown (Figure 7B). While there was no statistically significant difference in maximum rate of cleavage between NELF from males and females, there is a large degree of interindividual variability in rate of cleavage between donors.

**Fig 6.**
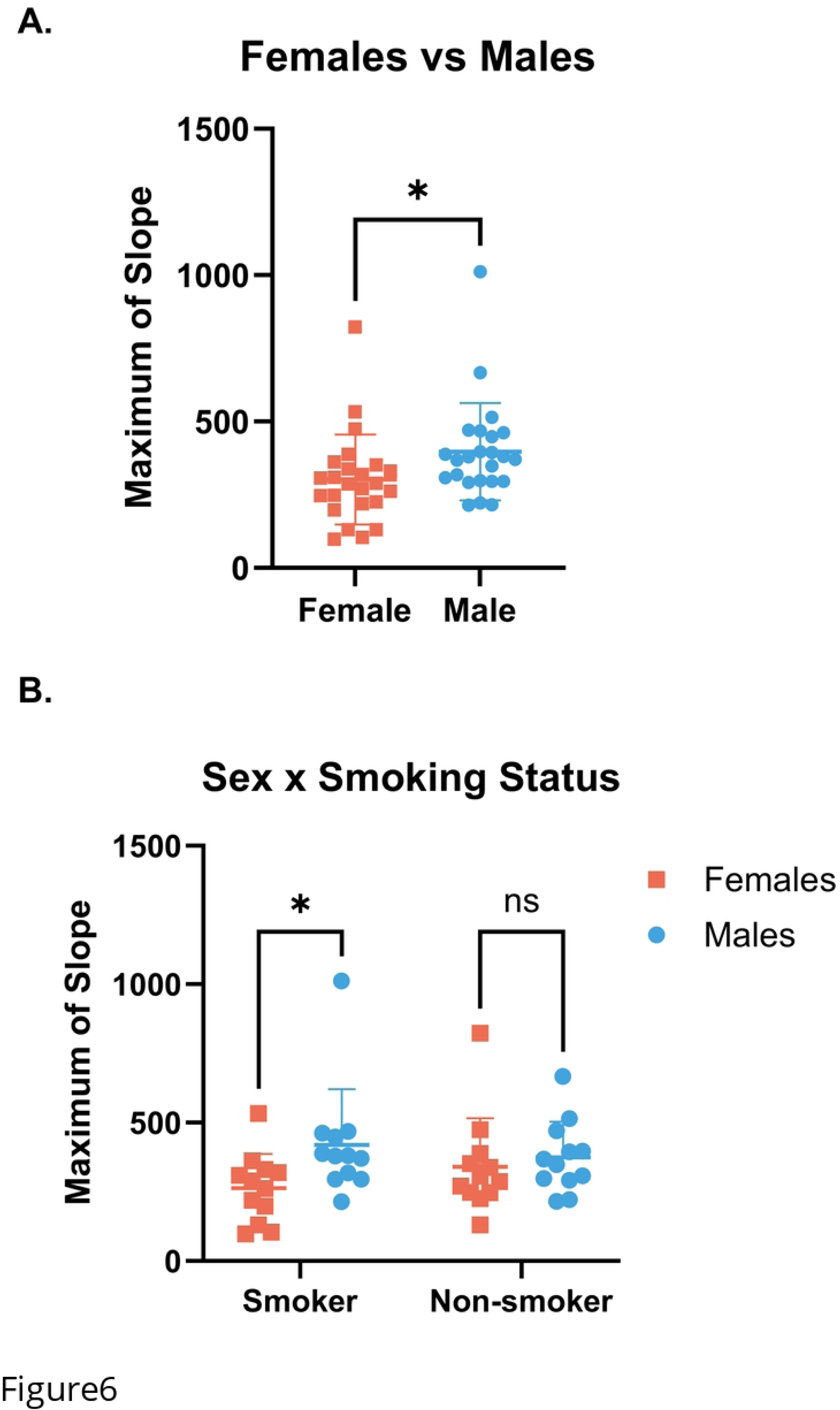
Differences in cleavage of Influenza H1 by proteases in NLF from male and female smokers. Rates of cleavage of the Influenza H1 peptide by NLF samples from smokers and non-smokers. No outliers were identified by the ROUT (Q=1%) method. A) Sex difference when the data are aggregated by smoking status (unpaired t-test, p≤0.05). B) Separating by smoking status demonstrates greater cleavage of the Influenza H1 peptide in samples from male donors (2-way ANOVA with Bonferroni‘s post hoc test, p≤0.05). In both plots, mean with standard deviation is shown.

**Fig 7.**
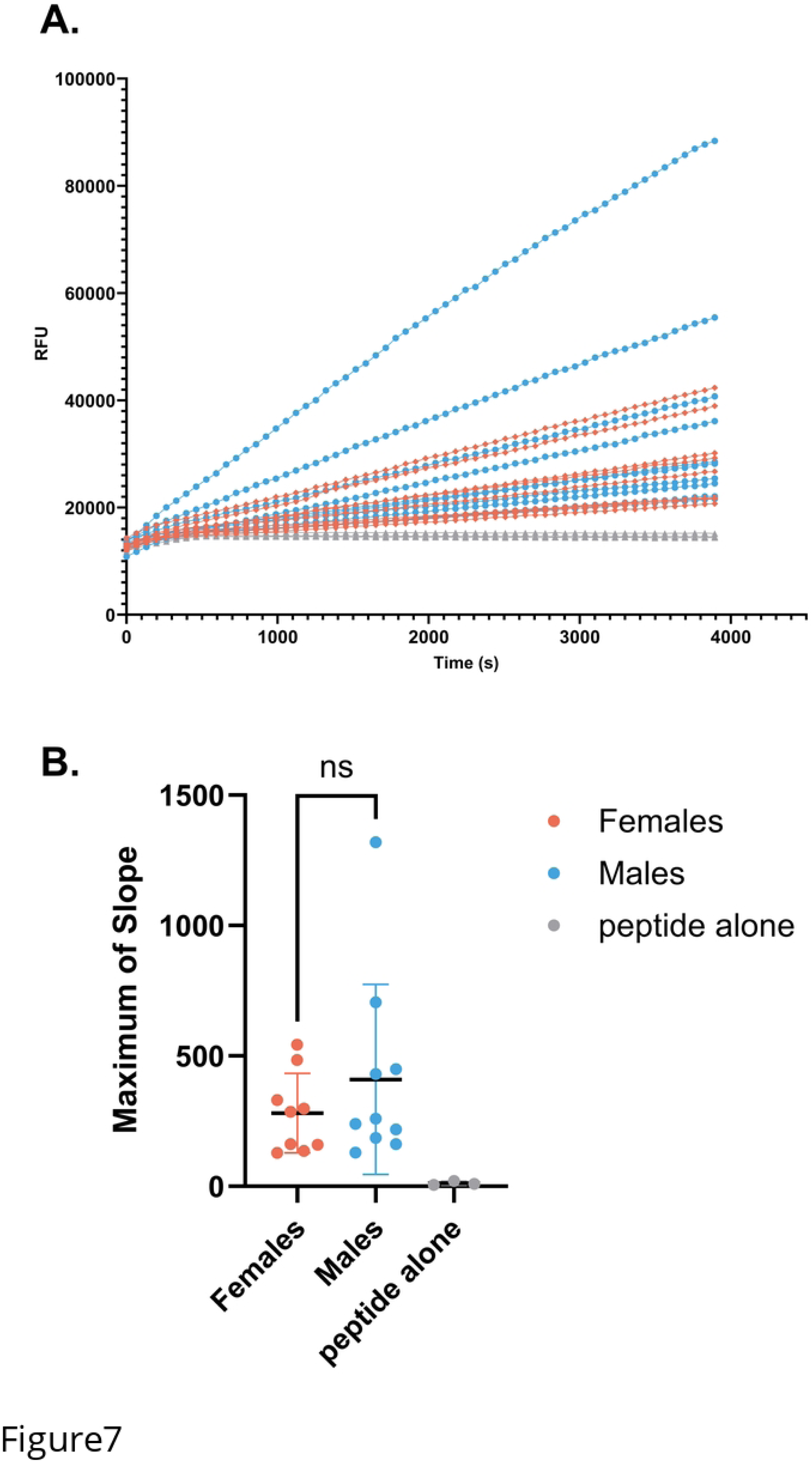
Proteases in NELF cleave Influenza H1. A) Cleavage of the Influenza H1 peptide by proteases in NELF samples from n=19 donors, males in blue (n=10), and females in orange (n=9). B) Maximum rate of cleavage for all samples. Maximums of slope were calculated for data in the range of time= 480-3894s. Welch’s unequal variance *t*-test was used to evaluate difference in maximum slope between sexes.

### ALI culture apical washes also cleave SARS-CoV S proteins

To demonstrate the utility of this assay to other viruses of public health relevance, we tested HNEC- and HBEC-mediated cleavage of the SARS-CoV peptides listed in Table 1. Figure 8A shows that enzymes in HNEC apical washes indeed cleave the SARS-CoV-2 S peptide, and addition of a protease inhibitor cocktail greatly decreases the maximum rate of cleavage of the peptide. Furthermore, addition of rhFurin to the SARS-CoV-2 S peptide reaction increases the rate of cleavage of this peptide (Figure 8B). In contrast, when rhFurin is added to the Influenza H1 cleavage reaction, there is a slight but statistically significant decrease in rate of peptide cleavage (Figure 8C). Furthermore, Figure 9A, B, and C demonstrate that proteases secreted from the apical surface of HNECs and HBECs grown at ALI successfully cleave the S peptides of SARS-CoV-1, SARS-CoV-2, and SARS-CoV-2 Delta, respectively. There was again no statistical difference in the maximum of slope between HNEC and HBEC cultures for any of the SARS-CoV peptides (Figure 9D). Interestingly, cleavage of the SARS-CoV-1 S peptide occurs at a much lower rate than for the other peptides. Figure 10 demonstrates proteases in NELF from the same donors used for the influenza H1 cleavage assay also cleave SARS-CoV-1, SARS-CoV-2, and SARS-CoV-2 Delta S peptides. Similar to influenza H1, there is much interindividual variability in cleavage of these peptides.

**Fig 8.**
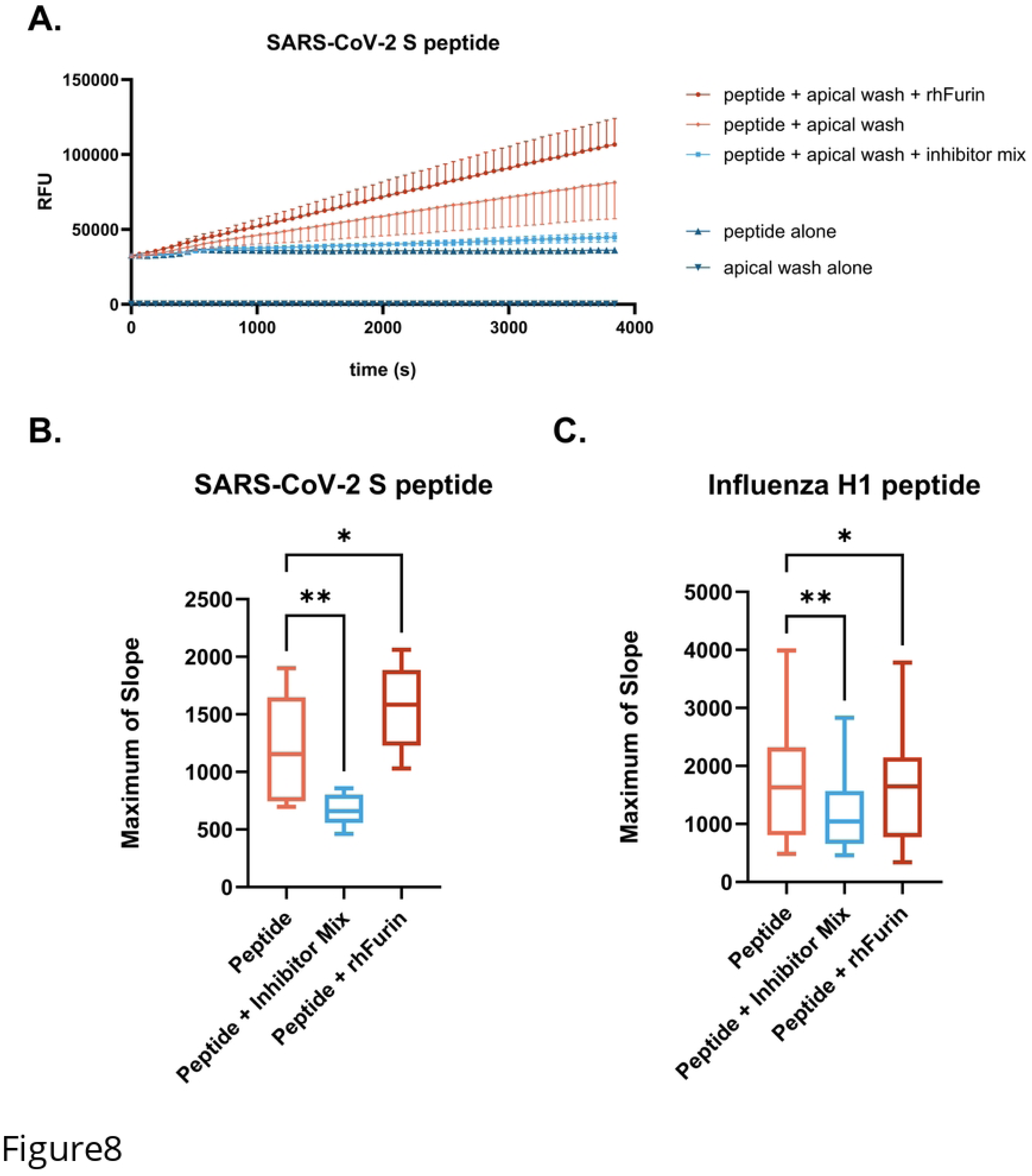
Cleavage of SARS-CoV-2 S IQF peptide by apical wash samples from n=9 HNEC donors. A) Change in RFU vs time. The addition of a mixture of protease inhibitors or rhFurin on rate of cleavage is also shown. Samples were collected 4 d since the prior apical wash. B) Maximum rates of reaction for SARS-CoV-2 S and C) for Influenza H1. Differences between groups were detected by repeated measures one-way ANOVA with Dunnett‘s post hoc test; * p≤0.05, ** p≤0.01.

**Fig 9.**
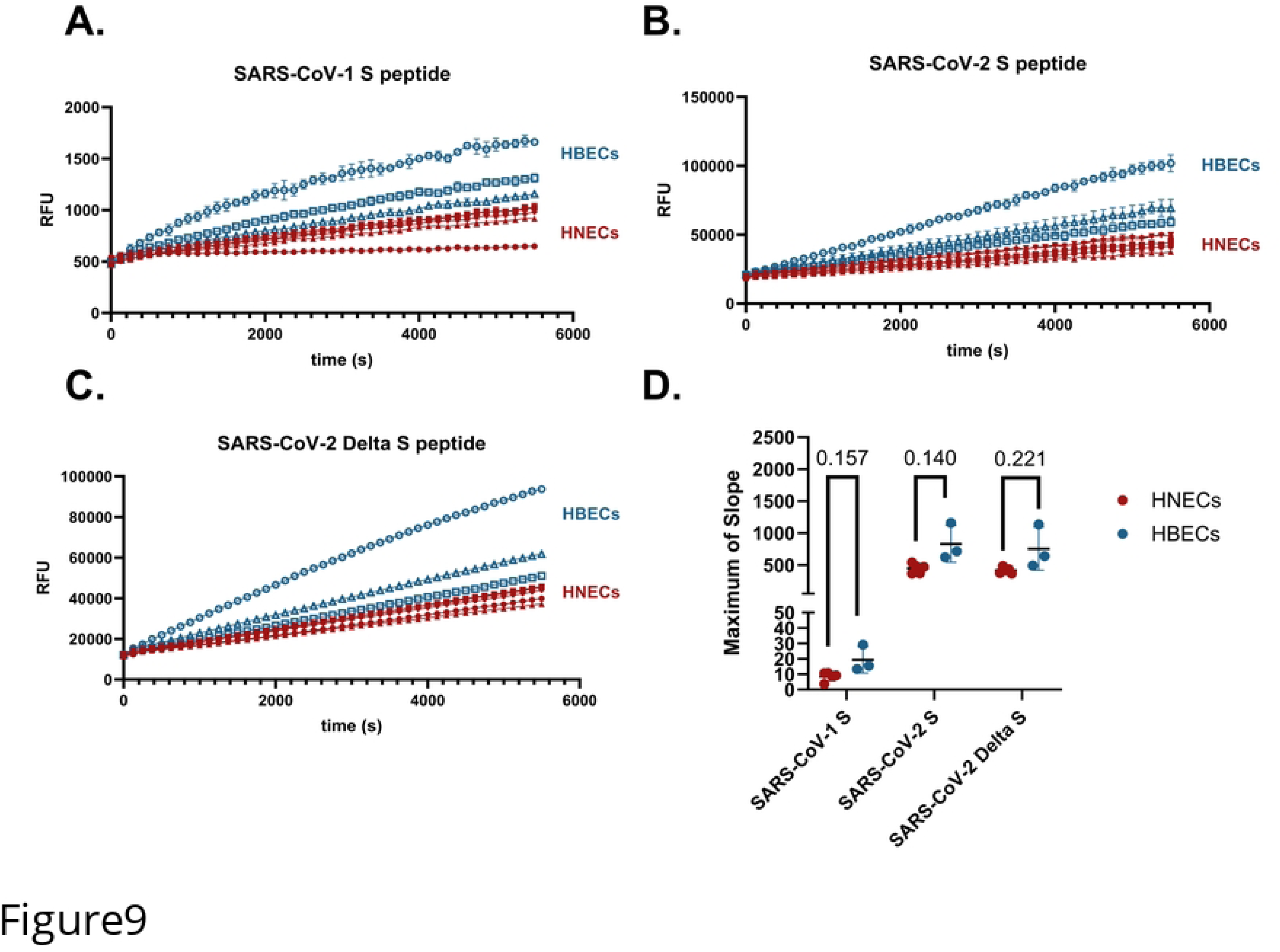
Proteases in HNEC and HBEC apical washes cleave SARS coronavirus peptides. Rates of cleavage of SARS S1/S2 IQF peptides by apical wash samples from n=5 HNEC donors and n=3 HBEC donors. Cleavage of A) SARS-CoV-1 B) SARS-CoV-2, and C) SARS-CoV-2 Delta. All samples were collected 7 d from the prior apical wash. Each line represents the mean of three technical replicates with standard deviation. D) Differences in maximum of slope between HBECs and HNECs for each peptide were tested with individual Welch’s unequal variance *t*-tests.

**Fig 10.**
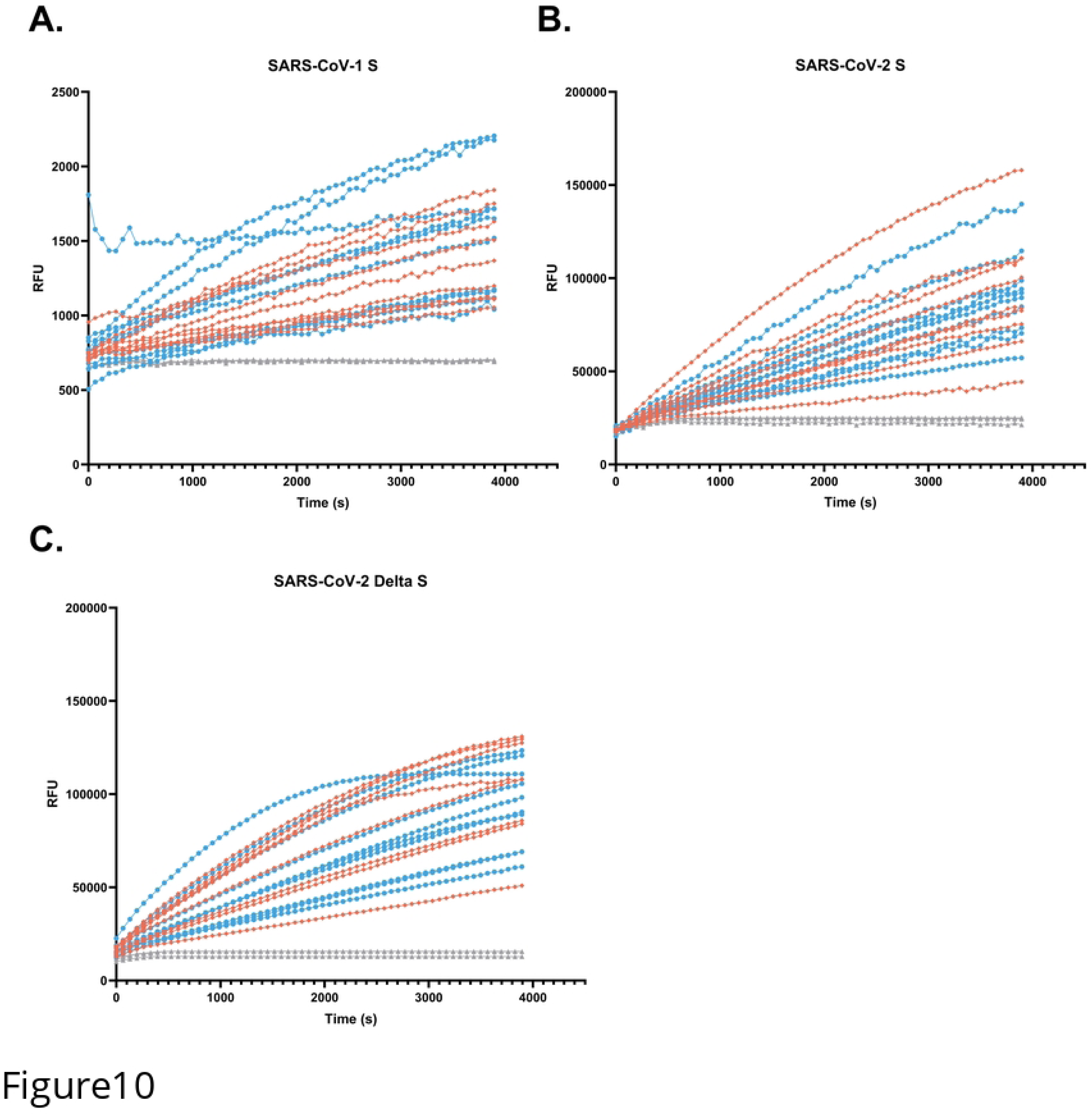
Proteases in NELF cleave SARS coronavirus peptides. Rate of cleavage of the SARS coronavirus peptides by proteases in NELF samples from n=19 donors, males in blue (n=10), and females in orange (n=9). Cleavage of SARS-CoV-1 is shown in (A), SARS-CoV-2 in (B) and SARS-CoV-2 Delta in (C).

## Discussion

We sought to better understand the proteolytic landscape present on the airway surface and demonstrate the utility of a simplified method for evaluating the overall proteolytic capacity of airway samples which does not rely on measurement of individual proteins. In airway surface liquid samples from primary *in vitro* airway cultures as well as clinical *ex vivo* samples, we identified numerous proteases and antiproteases with diverse but overlapping substrate specificities. Additionally, a method for efficiently probing proteolytic activity (in this case toward viral substrates) was presented, revealing active proteases can be easily measured in a variety of airway samples. Furthermore, these findings reveal interindividual differences in cleavage and activation of multiple airway viruses, possibly indicating differential susceptibility to infection. To demonstrate an application of the assay, cleavage of Influenza H1 peptide was compared in NLF samples from male and female smokers and non-smokers, revealing NLF from male smokers more readily cleaved Influenza H1 compared to female smokers.

The surfaces of the airways exemplify a complex proteolytic landscape populated with diverse and mechanistically redundant proteases and antiproteases, demonstrated in Tables 5 and 6. Previous studies have identified differentially expressed proteases or antiproteases in the proteomes of airway samples from healthy versus disease phenotypes (66, 67). However, to our knowledge, this study represents the first time specific proteomic profiling of both airway proteases and antiproteases has been assessed in airway samples from healthy donors. Our analysis puts a particular emphasis on proteases/antiproteases in the upper airway mucosa, which represents the initial target for many respiratory viruses. We identified 48 unique proteases in the HNEC apical wash samples and 57 in the NELF samples, which is in contrast to a former study which assessed the protease composition of bronchoalveolar lavage fluid from healthy donors and identified only 13 proteases (9). Moreover, this study compares the secreted proteases and antiproteases detected from *in vitro* primary nasal epithelial cells and *ex vivo* nasal epithelial lining fluid (NELF) from healthy human donors. Unsurprisingly, there were greater numbers of unique proteases and antiproteases detected in the NELF samples compared to the HNEC apical washes, which could be explained by the presence of cell types other than epithelial cells in the human nasal mucosa *in vivo*, including monocytes and neutrophils. However rather surprisingly, there were some proteases detected in HNEC apical wash samples which were not detected in NELF, including matriptase (ST14) and several metalloproteases (ADAM9, ADAM10, ADAM28). While this could reflect interindividual differences in nasal protease expression (since these samples were obtained from different donors), changes in cell biology induced from growing cells in culture could also provide an explanation. Although the data were not obtained from matched donors, we believe that the results are impactful in illustrating the complexity of the whole proteolytic environment found in this region of the airway. Further, these data emphasize the difficulty of predicting overall changes in the proteolytic environment of the airway based on changes in single enzymes or inhibitors, a method which has been previously used to predict changes in infection susceptibility.

Assaying proteolytic capacity of airway samples toward substrates of interest presents an effective method of detecting perturbations of the protease:antiprotease ratio following exposures or treatments and in the case of underlying diseases. The assay we show here offers advantages over methods employing infectious viruses and Western-blot to assess how changes in the proteolytic environment impact susceptibility to viral infection. Our data demonstrate that the assay developed and optimized here has utility in examining the proteolytic capacity of apical secretions from HNEC and HBEC organotypic cultures to activate multiple respiratory viruses of public health relevance (Figures 1, 4, 9, and 10). Additionally, because of their immediate human relevance, we demonstrate proteolytic activity in clinical human airway samples from healthy and exposed (i.e. smokers) populations analyzed *ex vivo*. We found that proteolytic activity can indeed be measured in stored nasal samples (Figures 6 and 7). These data show interindividual variability in proteolytic activity toward Influenza A virus, suggesting the assay’s utility in identifying potentially susceptible populations. Specifically, previous epidemiological studies have indicated female cigarette smokers have worse outcomes during Influenza infection than male smokers or non-smokers (68–70). Smoking has been found to increase expression of proteases such as neutrophil elastase, TMPRSS2, and certain matrix metalloproteases in the lungs (45, 71, 72). However, smoking also increases antiprotease SLPI expression (47). Thus, we wanted to evaluate whether NLF fluid from smokers and non-smokers would differentially cleave Influenza H1. We found no difference in cleavage of the Influenza H1 substrate in NLF samples from smokers and non-smokers. However, we report here a greater proteolytic activation of Influenza H1 in NLF from male smokers compared to female smokers (Figure 6), suggesting that factors other than initial infection of nasal epithelial cells may contribute to the greater severity of Influenza virus infections seen in female smokers.

Additionally, our data suggest that proteases accumulate on the apical surface of ALI cultures over time (Figure 2) and routine washing of these cultures removes proteases from the culture surface. This may be an important consideration for studies involving inoculation of the apical surface of cultures with respiratory viruses or evaluating efficacy of inhaled therapeutics. In addition, previous studies have suggested that only membrane-associated proteases are effective in cleaving Influenza H1 peptide and concluded secreted proteases are not active toward the virus (49, 50). However, our data do not support this finding as we observed proteolytic cleavage of Influenza H1 both in apical wash samples and directly on the cell surface.

With access to a plate reader with an atmospheric control unit, this assay can also be used to measure proteolytic cleavage directly on the apical surface of ALI cultures in real time, demonstrated in Figure 5. This expansion of our assay also allows the assessment of membrane-bound/tethered proteases in the respiratory mucosa. Dosing test compounds in the basolateral medium presents an application of the assay in testing efficacy of drugs designed for systemic circulation to modulate the protease:antiprotease balance on the airway surface.

The Spike protein of SARS coronaviruses undergoes two cleavage events to fuse with the host membrane; one at the S1/S2 site and another at the S2’ site (26). Similar to Influenza A, a variety of extracellular and membrane-bound host proteases cleave SARS-CoV-1 and SARS-CoV-2 at the S1/S2 site (37–39). In our assay, a mixture of chemical protease inhibitors against serine, cysteine, and Furin-like proteases reduced the cleavage rate of the SARS-CoV-2 peptide more dramatically than the Influenza H1 peptide in the same samples (Figure 8). This suggests that at least partially different subsets of proteases cleave these peptides. The increased rate of cleavage of the SARS-CoV-2 S peptide upon addition of rhFurin is expected since this virus acquired a Furin cleavage site in the S protein, characterized by multiple basic amino acids (typically Arginine) grouped together (underlined in Table 1) (73, 74). Influenza H1 does not contain a Furin cleavage site and addition of recombinant Furin slightly reduced the rate of cleavage of the Influenza H1 peptide. One possible explanation for this finding is competitive binding by the rhFurin to the peptide without catalysis of the cleavage reaction, reducing interactions between the peptide and proteases which successfully cleave it. Although the SARS-CoV-2 S1/S2 cleavage site has been previously shown to be cleaved by extracellular and membrane-bound proteases (37), it is likely that this site is primarily cleaved by intracellular Furin before progeny virions are released from an infected cell (26). Thus, modeling activation of this cleavage site by airway surface liquid samples is only an approximation for SARS-CoV-2 viral activation.

## Conclusions

The data shown here reinforce the vast diversity of unique proteases and antiproteases found in airway surface liquid samples from both organotypic airway culture models and clinical human samples. The methodology we describe is a straightforward, easy to use, and adaptable approach for measurement of the proteolytic activity of airway samples toward viral substrates, though the peptide design could be modified for any substrate of interest. This assay has utility for assessment of environmental exposures, disease states, or pharmaceutical interventions on the activity of airway proteases used by respiratory viruses to initiate infection. In addition, this methodology offers the ability to upscale to medium/high throughput and has greater practicality compared to measurement of cleavage rates by individual proteases or assessment of concentrations of individual proteases/antiproteases, which offer only an approximation of total proteolytic activity.

Supplemental Fig 1. Proteases remain enzymatically active with multiple freeze/thaws. Effects of 0-4 freeze and thaw cycles of apical wash samples from n=5 HNEC donors on proteolytic activity toward the Influenza H1 peptide. Lines represent individual donors and bars are means of all 5 biological replicates. There were no statistically significant differences between groups by repeated measures one-way ANOVA with Bonferroni’s post hoc test.

